# *Histophilus somni* Survives in Bovine Macrophages by Interfering with Phagosome-Lysosome Fusion, but Requires IbpA for Optimal Serum Resistance

**DOI:** 10.1101/322768

**Authors:** Yu Pan, Yuichi Tagawa, Anna Champion, Indra Sandal, Thomas J. Inzana

**Author notes:** Current Address: Memphis VA Medical Center, 1030 Jefferson Avenue, Memphis TN 38104 USA.

## Abstract

*Histophilus somni* survives intracellularly in professional phagocytic cells, but the mechanism of intracellular survival is not understood. The Fic motif within the DR1/DR2 IbpA fibrillar network protein of *H. somni* is cytotoxic to epithelial and phagocytic cells, which may interfere with the bactericidal activity of these cells. To determine the contribution of IbpA and Fic on resistance to host defenses, strains and mutants that lack all of or a small region of *ibpA* or DR1/DR2 were tested for survival in bovine monocytic cells and for serum susceptibility. A mutant lacking IbpA, but not DR1/DR2, was more susceptible to killing by antiserum than the parent. *H. somni* strains expressing IbpA replicated in bovine monocytes for at least 72 hours, and were toxic for these cells. Virulent strain 2336 with transposon insertions or deletions within IbpA remained toxic for bovine monocytes. However, strain 2336 mutants lacking all of *ibpA* or both DR1/DR2 were not toxic to the monocytes, but survived within the monocytes for at least 72 hours. Examination of intracellular trafficking of *H. somni* with monoclonal antibodies to early and late phagosomal markers indicated that early phagosomal marker EEA-1 colocalized with both disease isolate strain 2336 and serum-sensitve mucosal isolate strain 129Pt, but only strain 2336 did not co-localize with late lysosomal marker LAMP-2 and prevented acidification of phagosomes. These results indicate that virulent isolates of *H. somni* are capable of surviving within phagocytic cells through interference of phagosome-lysosome maturation. Therefore, *H. somni* may be considered a permissive intracellular pathogen.

---

The gram-negative bacterium *Histophilus somni* is an opportunistic pathogen associated with bovine respiratory disease and multi-systemic diseases in cattle and sometimes sheep, including thrombotic meningoencephalitis (TME), myocarditis, arthritis, mastitis, reproductive failure and abortion, and others; probably resulting from bacteremia (1). However, some strains of *H. somni* are serum-sensitive, and at least one such strain (129Pt) lacks many of the virulence factors associated with disease isolates (2). The only known reservoir for *H. somni* are the mucosal sites of ruminants (3).

Virulent strains of *H. somni* possess a wide variety of physiological properties and mechanisms that primarily protect the bacteria from host defenses or modulate host immune cells. Such mechanisms include phase variation of lipooligosaccharide (LOS), modification of LOS with sialic acid and phosphorylcholine (4), apoptosis of endothelial cells and neutrophils with disruption of intercellular junctions (5), and biofilm formation (6). Furthermore, the bacteria secrete a fibrillar and surface-associated immunoglobulin binding protein (IbpA), of which the *N*-terminus region is capable of binding immunoglobulins through their Fc component, and may also mediate adherence of the bacteria to host cells (7).The COOH-terminus of IbpA has homology to a region in YopT in *Yersinia* spp., but lacks cytotoxic activity (8). In contrast, sequence analysis of *ibpA* indicates that there are two direct repeats (DR1 and DR2) just upstream of the the *yopT*-like region, both of which contain a filamentation-induced by c-AMP (Fic) motif: HPFxxGNGR (8). These Fic domains can be found in both bacterial and eukaryotic cells. In *H. somni*, the Fic motifs of both DR1 and DR2 have been shown to be toxic for bovine epithelial and phagocytic cells, resulting in rounding up of the cells, increased detachment of infected macrophages, and disruption of actin fibers (9, 10). *H. somni* strain 2336 can inhibit phagocytosis of microspheres by primary bovine monocytes, but a mutant with essentially the entire *ibpA* gene deleted can not (9). Antibodies to the recombinant DR2 region of IbpA neutralize the cytotoxic effect on these cells (11). Immunization of mice and calves with recombinant DR2 also protects the animals from *H. somni* bacteremia and pneumonia, respectively (12, 13). The presence of IbpA on *H. somni* strains is also associated with serum resistance (7).

Virulent strains of *H. somni* are capable of surviving within bovine polymorphonuclear leukocytes (PMNs), monocytes, and macrophages (14, 15). Phagocytic cells infected with live *H. somni* are less capable of internalizing a secondary target, such as opsonized *Staphylococcus aureus* and microspheres (16, 17). Killed, whole bacteria or supernatant from heat-killed bacteria can also inhibit the internalization of *S. aureus* by PMNs, but not bovine macrophages (16, 17). We have previously reported that the oxidative burst generated by phagocytic cells in contact with viable disease isolates of *H. somni* is significanlty inhibited. However, there is no inhibition of the oxidative burst by killed *H. somni*, nonvirulent mucosal strain 129Pt, and heterologous strains that include *Haemophilus influenzae* and *Brucella abortus* (18). The mechanism by which *H. somni* survives within phagocytic cells remains unclear. Because the Fic motifs within IbpA are toxic to phagocytic cells and induce disruption of actin filaments, it is possible that *H. somni* survives intracellular killing through Fic-mediated interference of phagocytotic cell functions. In this study, we used various mutants with transposon (Tn) insertions and in-frame deletions in *ibpA* to examine the contribution of IbpA and the Fic motifs to serum susceptibility and intracellular killing of *H. somni*, and how virulent disease isolates and avirulent isolates traffic within bovine monocytes.

## RESULTS

### Intracellular survival of *H. somni* in bovine monocyte (BM) and bovine peripheral blood monocyte cells (BPBM)

Several macrophage or monocyte cell lines, including BM, bovine FBM-17, mouse J774A.1, and human THP cells, were examined for the capability of *H. somni* strain 2336 to survive intracellularly in comparison to freshly collected BPBM (data not shown). The BM cell line was found to be most comparable to BPBM in regard to intracellular survival or killing of *H. somni. H. somni* pathogenic strain 2336 survived in BPBM and was cytotoxic to these cells resulting in detachment and rounding up of the cells (data not shown), as previously described for FBM-17 cells (9). In contrast mucosal strain 129Pt from the healthy prepuce was not cytotoxic and did not survive in BM cells (Fig. 1). Strain 2336 was also capable of replicating in the BM cell line, but strain 129Pt was not, which was similar to the results obtained with PMBC (Fig. 1). Mouse macrophage cell line J774.1 and human leukemic cell line THP-1 were also tested, but were unable to kill strain 129Pt (data not shown). Therefore, BM cells were used for all subsequent studies.

**Figure 1.**
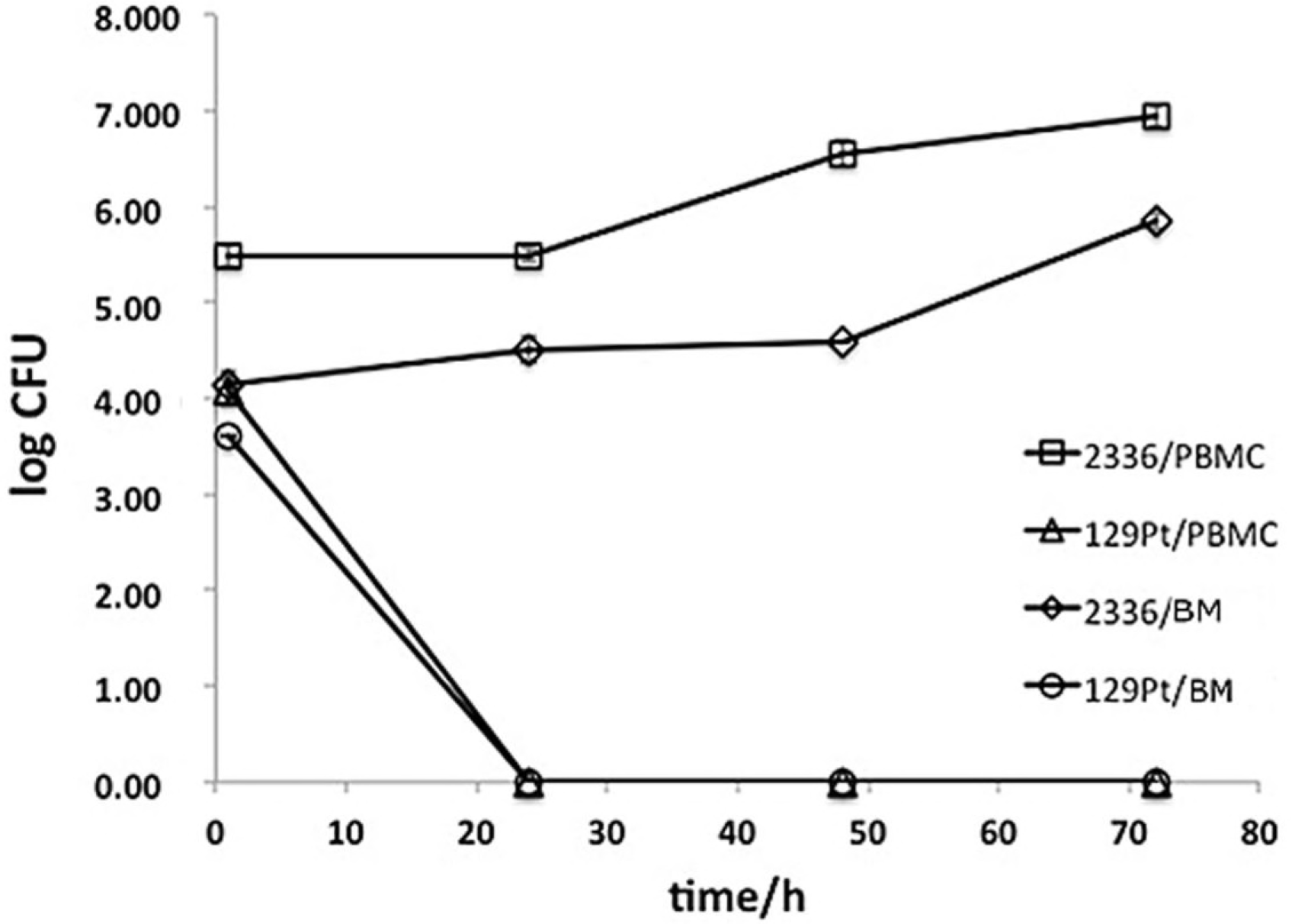
Survival of *H. somni* strains 2336 and 129Pt in BPBM and BM monocyte cell line. The bacteria were incubated with monocyte cells for 1 hr (uptake time) at 100:1 (bacteria:monocytes), gentamicin added to kill extracellular bacteria, and after 30 min the moncytes were washed, incubated for 0, 24, 48, or 72 hr, lysed, and the released bacteria cultured on blood agar. □, strain 2336 incubated with BPBM; Δ, strain 129Pt incubated with BPBM; ◊, strain 2336 incubated with BM cells; ○, strain 129Pt incubated with BM cells. Results represent the mean ± standard deviation of at least 3 experiments. Standard deviation bars are difficult to see due to their small size and the symbols.

Several *H. somni* strains from disease sites and healthy mucosal sites were tested for their ability to survive within BM cells following phagocytosis. The intracellular number of all isolates from disease sites and some isolates from healthy genital sites increased after 24 h co-incubation with the monocytes (Fig. 2). The Fic domains within DR1/DR2 of IbpA have been shown to be cytotoxic to host cells (9), and the presence or absence of *ibpA* in all the *H. somni* strains used in this study has been previously documented (Table 1) (19). All the disease isolates and most of the vaginal isolatess tested were able to replicate intracellularly, but most preputial isolates tested did not (Table 1 and Fig. 2). Strains 1P, 129Pt, 130Pf, and 133P from the bovne prepuce do not produce IbpA (19) and were unable to replicate intracellularly after 24 h co-incubation (Table 1 and Fig. 2). However, some preputial isolates previously shown to produce IbpA (24P, 124P, and 20P) were also killed by BM cells, though preputial IbpA-producing strain 22P was resistant. Of interest was that strain 1225, which was isolated from the bovine prepuce in The Netherlands, was highly resistant to intracellular killing, but it was unknown whether this isolate was associated with disease or expressed IbpA. Therefore, the expresion of IbpA was not universally associated with intracellular survival.

**Figure 2.**
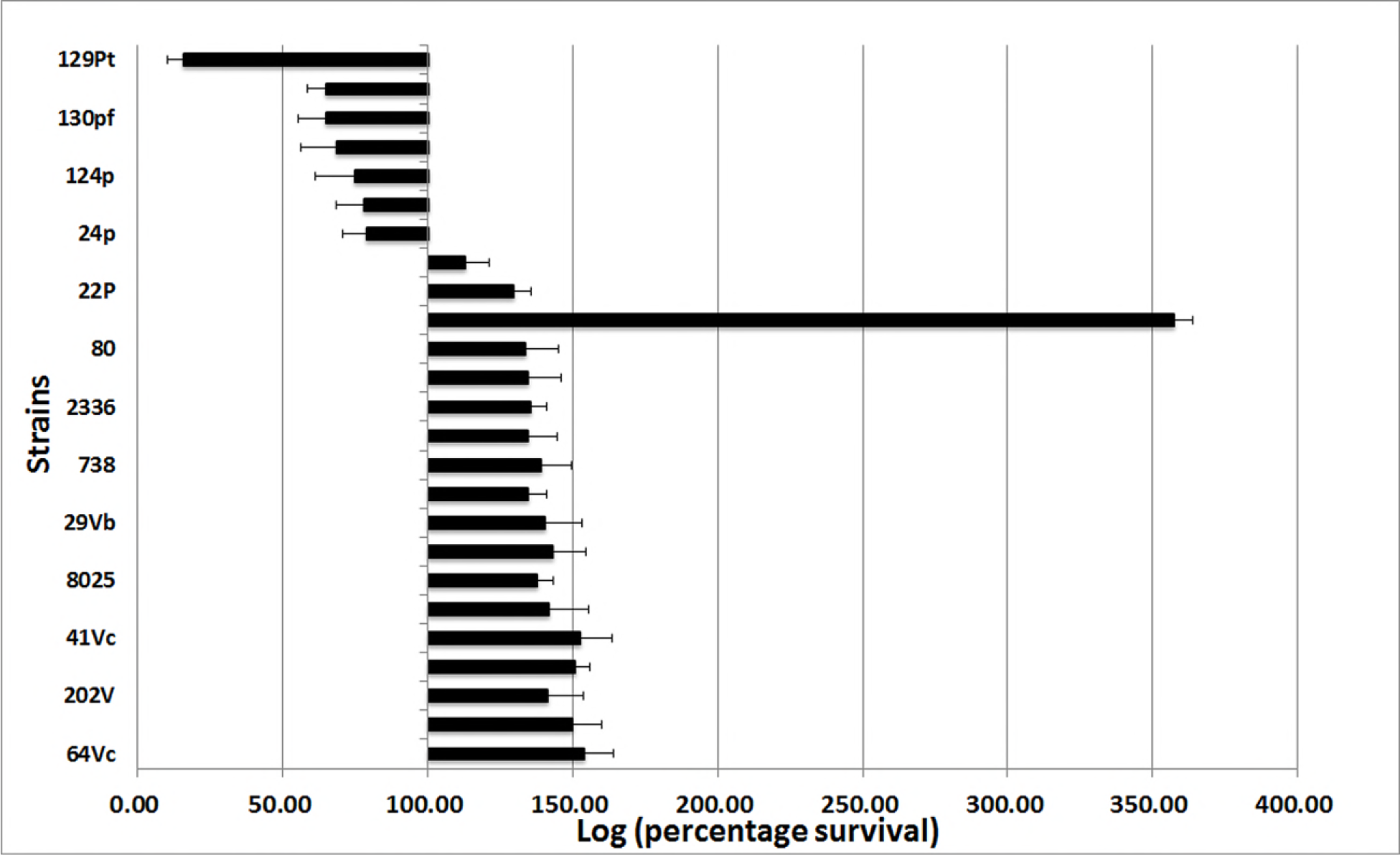
Survival of *H. somni* strains in BM monocytes after 24 h co-incubation. The strains on the left of 100% were killed in the monocytes; the strains on the right of 100% replicated in the BM cells. Percent survival of *H. somni* was determined by lysis of BM cells after 1 h (incubation time given for uptake) and 24 h post co-incubation, and culture. The number of colonies recovered after 24 h incubation was divided by the number of colonies after 1 h of incubation × 100. Results represent the mean ± standard deviation of 3 experiments.

**Table 1:** *H. somni* trains used in this study.

| 581 | Strain | Description | IbpA present<sup>a</sup> | Serum resistant | Reference or source |
| --- | --- | --- | --- | --- | --- |
| 582 | 2336<sup>b</sup> | Isolated from bovine pneumonia strain |  | ++ | (26) |
| 583 | 2336::*TnfhaB* #3 | *ibpA* Tn mutant | + | ND<sup>c</sup> | (29) |
| 584 | 2336::*TnfhaB* #13 | *ibpA* Tn mutant | + | ND | (29) |
| 585 | 2336::*TnfhaB* #91 | *ibpA* Tn mutant | + | ND | (29) |
| 586 | 2336::*TnfhaB* #9 | *ibpA* Tn mutant | + | ND<sup>b</sup> | (29) |
| 587 | 2336::*TnfhaB* #27 | *ibpA* Tn mutant | + | ND | (29) |
| 588 | 2336::*TnfhaB* #23 | *ibpA* Tn mutant | + | ND | (29) |
| 589 | 2336::*TnfhaB* #137 | *ibpA* Tn mutant | + | ND | (29) |
| 590 | 2336Δ*ibpA1* | *ibpA* deletion mutant (all of *ibpA*) |  | - | (9) |
| 591 | 2336Δ*ibpA5* | *ibpA* deletion mutant (3'-terminal sequence) |  | ND | (38, 39) |
| 592 | 2336Δ*ibpA7* | *ibpA* deletion mutant (DR2 sequence) |  | ND | (38, 39) |
| 593 | 2336Δ*ibpA8* | *ibpA* deletion mutant (DR1 sequence) | | ND | (38, 39) |
| 594 | 2336Δ*ibpA9* | *ibpA* deletion mutant (DR1 andDR2 sequence; | | | |
| 595 |  | all of Fic motif) |  | + | (38, 39) |
| 596 | 2336Δ*ibpA11* | *ibpA* deletion mutant (R1R2 sequence) | | ND | (38, 39) |
| 597 | 129Pt | normal prepuce | - | - | (26) |
| 598 | 1P | normal prepuce | - | - | (26) |
| 599 | 130Pf | normal prepuce | - | - | (26) |
| 600 | 133P | normal prepuce | - | - | (26) |
| 601 | 124P | normal prepuce | + | - | (26) |
| 602 | 20P | normal prepuce | + | + | (26) |
| 603 | 24P | normal prepuce | + | - | (26) |
| 604 | 221V | normal vagina | + | - | (26) |
| 605 | 22P | normal prepuce | + | + | (26) |
| 606 | 1225 | prepuce | ND | ND | V. Fussing, Natl. Vet. |
| 607 |  |  |  |  | Laboratry, Denmark |
| 608 | 80 | normal vagina | + | + | (26) |
| 609 | 1297 | pneumonia | + | + | (26) |
| 610 | 5166 | pneumonia | ND | ND | (26) |
| 611 | 738 | phase variant of 2336 |  |  |  |
| 612 |  | isolated from challenged calf lung | + | + | (46) |
| 613 | 649 | abortion | + | + | (26) |
| 614 | 29Vb | normal vagina | + | - | (26) |
| 615 | 318 | phase variat of strain 738 | + | + | (26) |
| 616 | 8025 | TME | + | + | (26) |
| 617 | 2089 | abortion | + | + | (26) |
| 618 | 41Vc | normal vagina | + | - | (26) |
| 619 | 208V | normal vagina | + | - | (26) |
| 620 | 202V | normal vaginal | + | - | (26) |
| 621 | 570 | abortion | + | - | (26) |
| 622 | 64Vc | normal vagina | + | + | (26) |

<sup>a</sup>The presence or absence of *ibpA* was determined by PCR and the production of IbpA by Western Blotting of the bacteria culture supernatant (19).
<sup>b</sup>All mutants were derived from strain 2336.
<sup>c</sup>ND – not determined.

### The role of IbpA in intracellular survival of *H. somni*

To assess the direct effect of IbpA on *H. somni* intracellular survival, BM cells were incubated for 2 hrs with *H. somni* culture supernatant concentrated 1:4 or 1:20 to enrich for IbpA, then infected with *H. somni* strain 129Pt, which cannot survive intracellularly. After 24 h incubation the intracellular survival of strain 129Pt in BM cells incubated with culture supernatant concentrated 1:4 or 1:20 was significantly (*p* < 0.001) greater than for the bacteria within monocytes not incubated with IbpA, and this effect was dose-dependent (Fig. 3). These results indicated that IbpA had a negative effect on BM cells to subsequently take up and/or kill strain 129Pt. IbpA (20-fold concentrate) also significantly enhanced the intracellular survival of the other *H. somni* strains (24 h incubation in monocytes) that were not as susceptible to intracellular killing as strain 129Pt, but also lacked the *ibpA* gene (1P, 133P, and 130Pf; *p* = 0.004, 0.001, and 0.005, respectively), although not to the extent as for strain 129Pt (Fig. 4). However, the addition of IbpA only moderately enhanced intracellular survival of strain 24P and, had no effect on strains 124P and 20P, (Fig. 4), all of which, as expected, produce IbpA (19).

**Figure 3.**
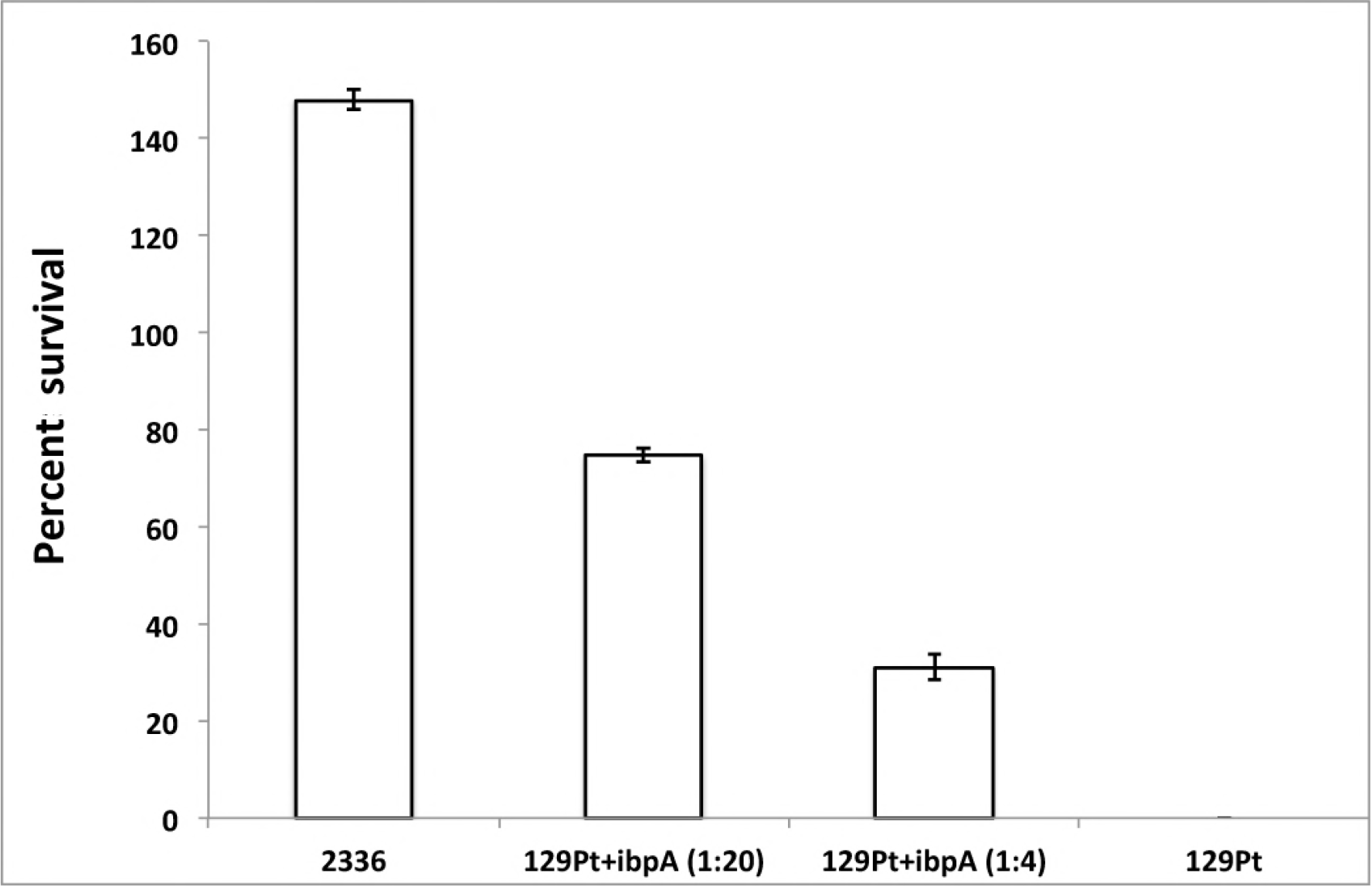
Percent intracellular survival of *H. somni* 129Pt after 24 h incubation in BM cells preincubated for 2 hr with or without 20-fold or 4-fold concentrated, semi-purified IbpA from 2336 supernatant. Strain 2336 survival shown for comparison. Results represent the mean ± standard deviation of at least 3 experiments.

**Figure 4.**
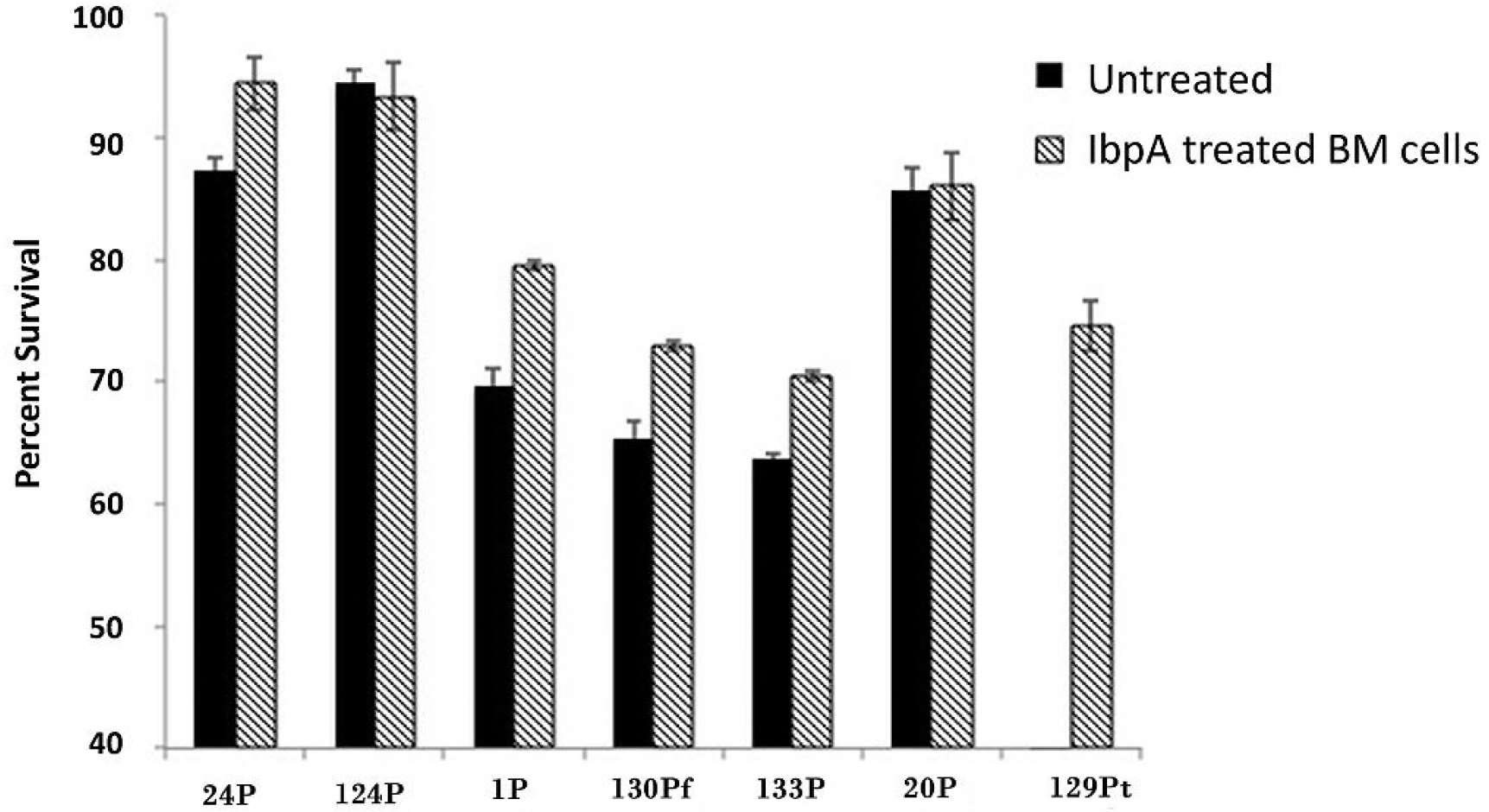
Percent survival of *H.somni* preputial isolates after 24 h incubation in BM cells that have been preincubated for 2 h with 20-fold concentrated culture supernatant containing IbpA. Strains 129Pt, 1P, 133P, and 130Pf lack IbpA and were significantly more resistant to intracellular killing by BM monocytes after addition of IbpA. Strains 24P, and 124P and 20P produce IbpA and there was little or no difference, respectively, in killing by BM cells following preincubation of the cells with IbpA. Results represent the mean ± standard deviation of at least 3 experiments.

### Survival of *H. somni ibpA* mutants in BM cells

Several mutants with Tn insertion mutations in *ibpA* were selected from a bank of Tn mutants for intracellular survivial in BM cells. All the mutants replicated significantly more slowly than the parent strain at 24 h post-incubation with BM cells (*p* < 0.001), but none of the mutants demonstrated a significant difference in the number of viable intracellular bacteria after 48 or 72 h of incubation (*p* > 0.05) (data not shown). However, all of these mutants contained the Tn in the region encoding for Fc binding by IbpA, and none contained the tranposon within the DR1/DR2 region. Mutants with the *ibpA* gene replaced with a Kn^R^ gene or with in-frame deletions in *ibpA* were also tested for intracellular survival (Fig. 5). Mutants with essentially the entire *ibpA* gene removed (2336ΔIbpA1) or both DR1 and DR2 (DR1/DR2) containing the Fic motifs (2336ΔIbpA9) were capable of surviving within BM cells as effectively as parent strain 2336. Strains 2336ΔIbpA5 (deletion near 3’-terminus), 2336ΔA7 (DR2 only deleted), 2336ΔA8 (DR1 only deleted), and 2336ΔA11 (deletion of R1R2 sequences) were also no more susceptible to intracellular killing than the parent (data not shown). Therefore, destruction of actin filaments and cell toxicity due to the Fic motifs were not, by themselves, the mechanism by which *H. somni* survived within phagocytic cells. Furthermore, there was not a significant difference between the presence or absence of IbpA, DR1/DR2, and uptake of the bacteria.

**Figure 5.**
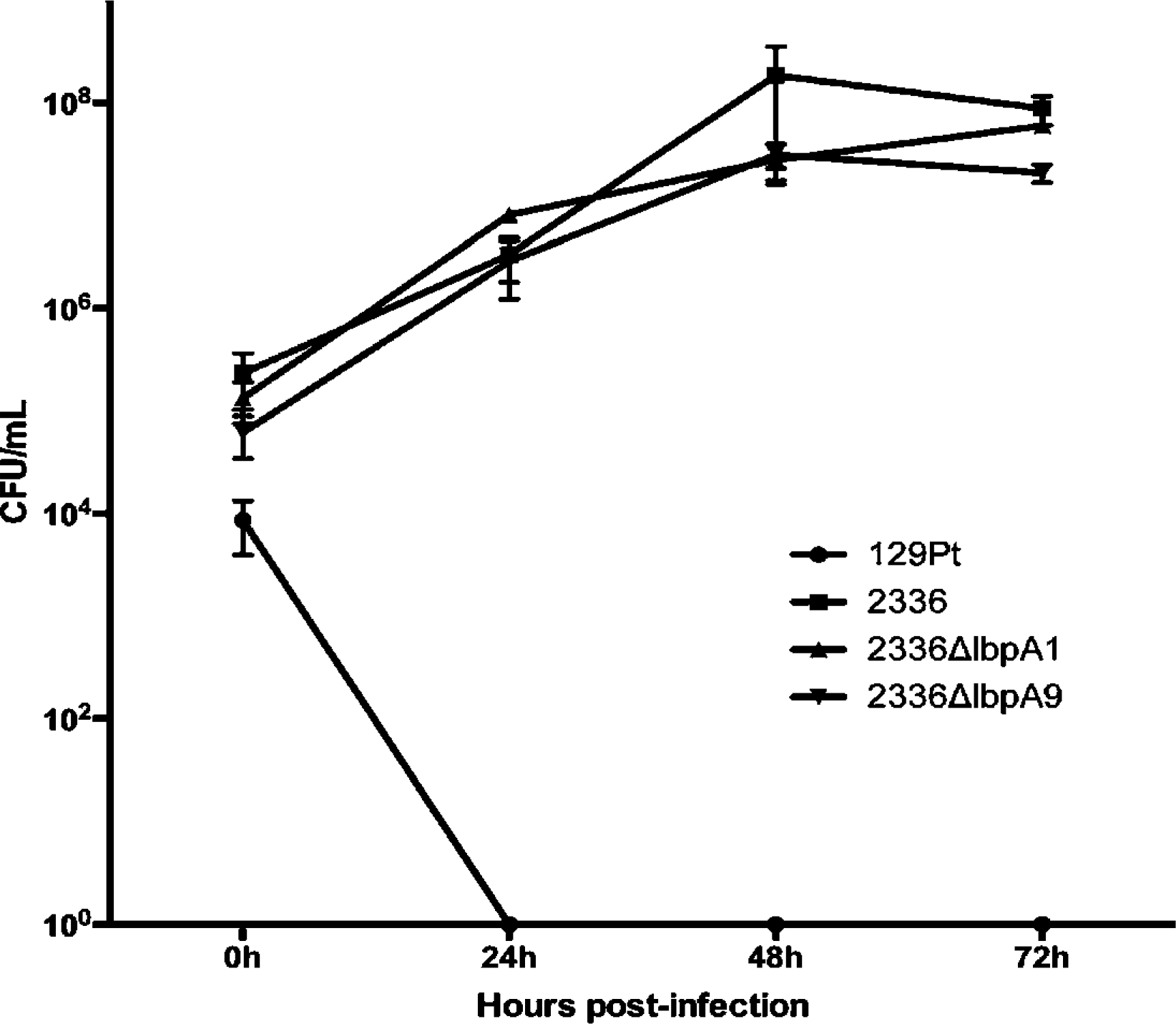
Intracellular survival of *H. somni ibpA* mutants and control strains in BM cells over 72 hr. Survival of strain 129Pt (•); strain 2336 (▪); strain 2336ΔIbpA1 (▲); strain 2336ΔIbpA9 (♦). The lack of DR1/DR2 repeats containing the toxic Fic motifs or essentially the entire IbpA protein had no effect on itracellular survival of strain 2336 in BM cells. Results represent the mean ± standard deviation of at least 3 experiments.

### Intracellular trafficking of *H. somni* within BM cells

Intracellular trafficking of pneumonia isolate strain 2336 in comparison to mucosal isolate strain 129Pt was examined by confocal microscopy to further clarify the mechanism of *H. somni* intracellular survival. Early phagosomal marker EEA-1 and late lysosomal marker LAMP-2 were both expressed in BM and co-localized with strain 129Pt with and without the addition of IbpA, and with strain 2336. Phagolysosomes containing strain 129Pt with and without the addition of IbpA were also acidified and co-localized with LAMP-2 (Figs. 6 and 7, *p* > 0.05). Therefore, intracellular trafficking of strain 129Pt in BM was not affected by the addition of IbpA. However, although EEA-1 co-locallized with strain 2336, the acidification of phagolysosomes and the expression/co-localization of LAMP-2 in BM cells infected with strain 2336 was significantly lower than with strain 129Pt (Figs. 6 and 7, *p* = 0.008).

**Figure 6.**
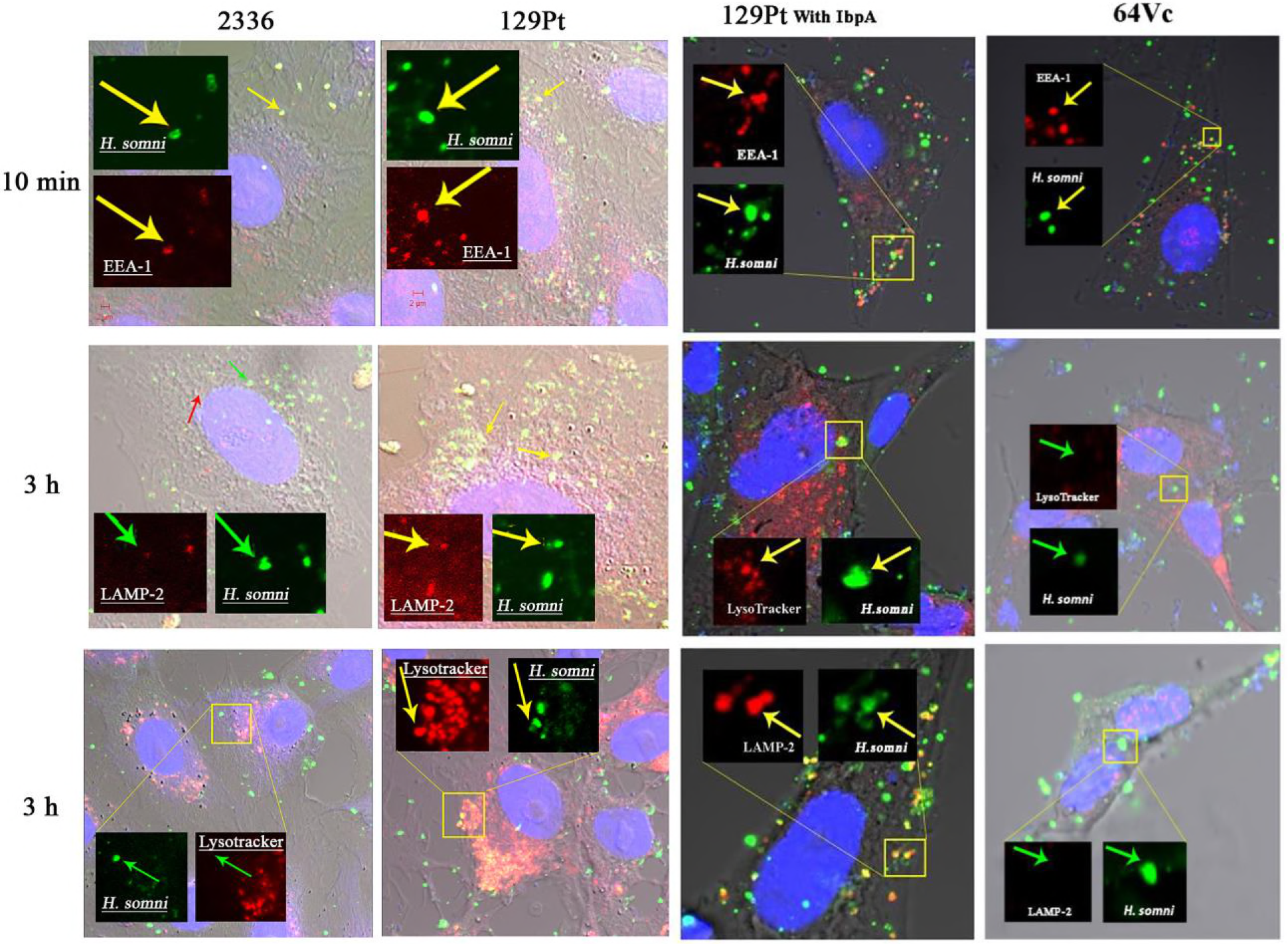
Confocal microscopy of *H. somni* strains 2336, strain 129Pt with and without addition of IbpA, and 64Vc following phagocytosis. The markers EEA-1 (10 min), and LAMP-2 and Lysotracker (3 hr) were stained with Alexa Fluor^®^ 546 (red) and *H. somni* with Alexa Fluor^®^ 488 (green). The red arrowhead point to the markers, the green arrowheads point to *H. somni*, the yellow arrowheads point to colocalization of the markers with *H. somni*. Each photo shown is a representative field of 6-10 fields examined from 3 separate experiments.

**Figure 7.**
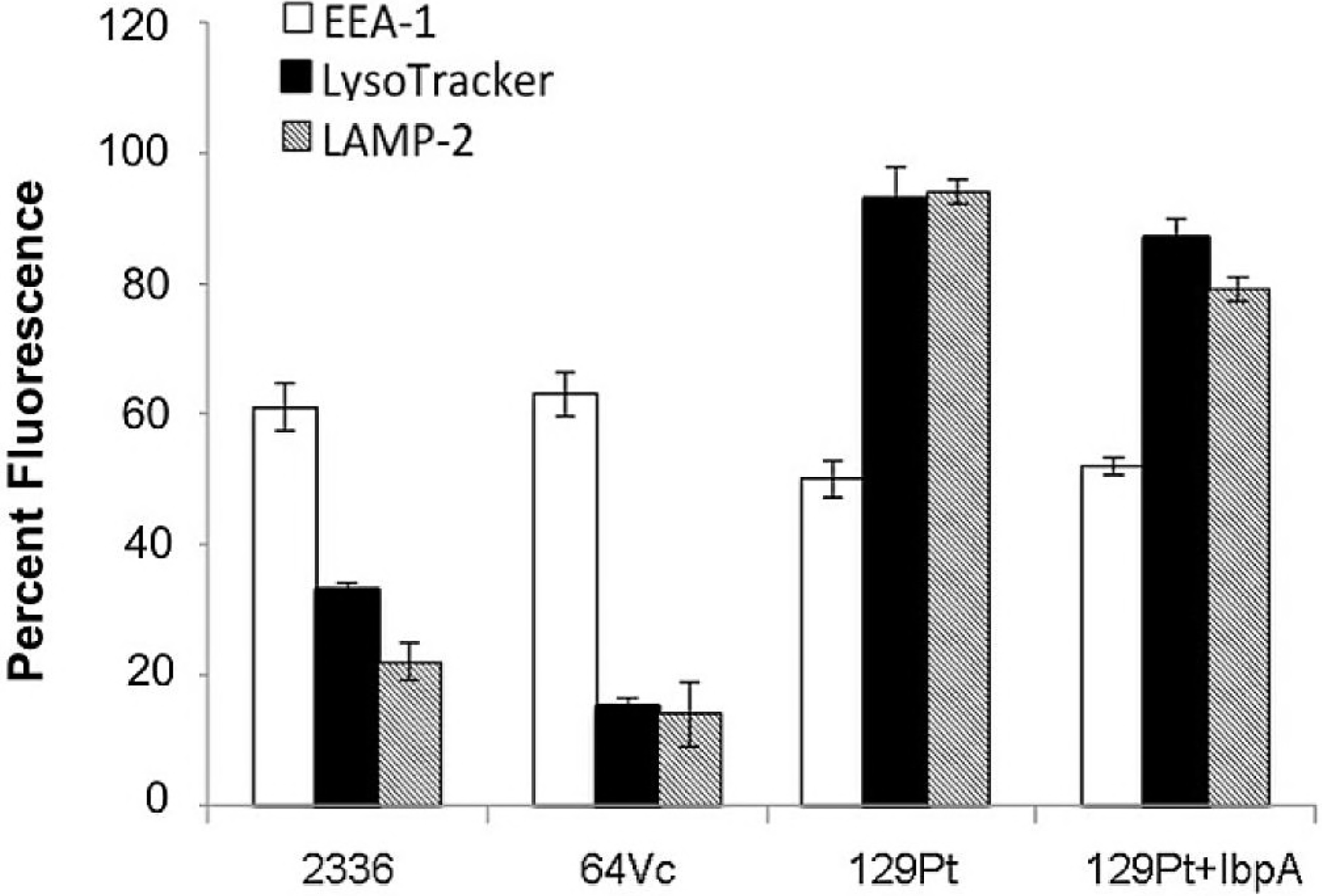
Quantification of *H. somni* serum resistant strains 2336 and 64Vc, and serum sensitive strain 129Pt (with and without added IbpA) that co-localized with markers EEA-1 (white column), LysoTracker (black column) or LAMP-2 (gray column). The co-localization of *H. somni* with each marker was quantified at different time points: EEA-1 (10 min), LysoTracker/acidification and LAMP-2 (3 h). Quantification of EEA-1 and LAMP-2 markers, and LysoTracker, was determined using the software imageJ and MicrobeTracker, which analyze the percentage of fluorescent dots and fluorescence signals, respectively. Results represent the mean ± standard deviation of at least 3 experiments.

The vaginal isolates used in this study were also capable of surviving in BM, although these isolates were not associated with disease. To compare the intracellular trafficking of a typical vaginal isolate to disease isolate strain 2336, strain 64Vc was also examined by confocal microscopy. Strain 64Vc co-localized with early phagosomal marker EEA-1 10 min after infection, but also inhibited the expression of LAMP-2 (*p* = 0.01) and the acidification of the phagosomes (*p* = 0.001 compared to 129Pt) (Figs. 6 and 7).

### Serum susceptibility of IbpA and DR1/DR2 mutants

*Histophilus somni* expression of IbpA is also asociated with serum resistance (7). Therefore, we sought to determine if a normally serum resistant strain becomes serum sensitive with the loss of IbpA or the DR1/DR2 regions. Serum resistance is relative in *H. somni*; even “serum resistant” strains can be killed in the presence of adequate antibodies to *H. somni* surface antigens. Therefore, to maximize the serum bactericidal effect, antiserum to *H. somni* LOS was used in the presence of pre-colostral calf serum as an antibody-free source of complement. In the absence of any antiserum, strain 2336 and mutants lacking essentially all of IbpA (*H. somni*ΔIbpA1) or both DR1 and DR2 (*H. somni*ΔIbpA9) increased in numbers, indicating these bacteria were resistant to the effects of complement alone (Fig. 8). At 40% antiserum, all the strains were effectively killed. However, in the presence of between 10% and 30% antiserum mutant *H. somni*Δ2336IbpA9, lacking DR1/DR2, was significantly more resistant to killing than even strain 2336 (*p* <0.008), but *H. somni* mutant 2336ΔIbpA1 (lacking all of IbpA) was more susceptible to killing in 10% antiserum (*p* = 0.004) than strain 2336, though not as susceptible as strain 129Pt. Therefore, the IbpA protein, but not the DR1/DR2 repeats containing the Fic motifs, contributed to serum resistance. However, other factors that may also be deficient in strain 129Pt appear to contribute to serum resistance in *H. somni*.

**Figure 8.**
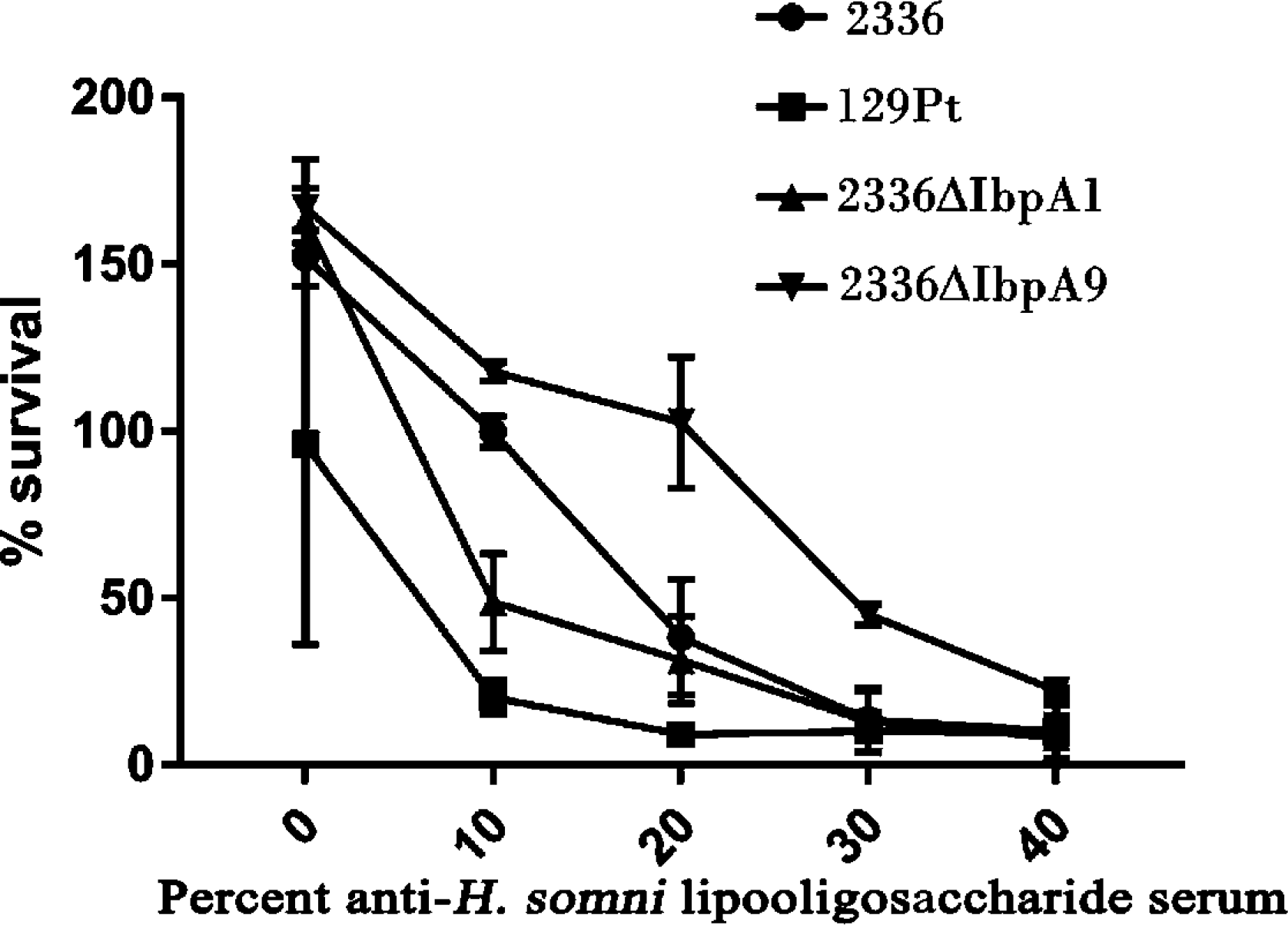
Serum resistance or susceptibiilty of *H. somni* and *ibpA* mutants. Isogenic mutants of control strain 2336 (•; serum resistant) lacking all of *ibpA* (▴; 2336ΔIbpA1) or only DR1/DR2 repeats containing the toxic Fic motifs (▾; 2336ΔIbpA9) were tested for susceptibility to killing by antiserum to *H. somni* lipooligosaccharide and bovine complement in comparison to control strain 129Pt (▪; serum sensitive). Results represent the mean ± standard deviation of at least 3 experiments.

## DISCUSSION

Virulent strains of *H. somni* are readily phagocytosed, but not killed, by neutrophils, macrophages, or monocytic cells (14, 15, 20). However, the mechanism by which *H. somni* surives within these cells is not clear. Generation of reactive oxygen intermediates are an important defense mechanism that phagocytic cells use to kill bacteria following phagocytosis (21). Inhibition of the oxidative burst by phagocytic cells following incubation with *H. somni* has been well established and may contribute to intracellular survival (14, 17, 18, 22–24). Several investigators have shown that this inhibition of the oxidative burst requires contact with, or the presence of, viable *H. somni* (16, 18, 22, 24), whereas others have reported such inhibitory activity can occur by killed cells or cell fractions (17, 25). The reason for this difference is unclear, but may be related to differences in the assays used. Furthermore, while disease isolates of *H. somni* are very efficient at inhibiting the oxidative burst of phagocytic cells, serum-sensitive isolates from the normal bovine prepuce are not (18). In addition, incubation of phagocytic cells with *H. somni* inhibits their subsequent uptake of opsonized *S. aureus* (16, 17, 22), indicating that the cells have been compromised in regard to phagocytic capacity following incubation with *H. somni*.

Bovine BM cells killed strain 129Pt as effectively as BPBM, but bovine FBM-17 cells, murine J774A.1, and human THP cells did not, indicating that the BM cells are well adapted to *H. somni* or that some immortalized cell lines may be deficient in aspects of intracellular killing. Strain 2336 survived within BM cells as well as in BPBM, indicating this cell line was a suitable surogate for BPBM for use in these assays. In this study, additional disease and mucosal isolates were tested for intracellular survival. However, while most preputial isolates were confirmed to be less capable of replicating in monocytes, isolates from the healthy vagina were as capable of surviving intracellularly as disease isolates. Of interest was that the isolates resistant to intracellular killing are also serum-resistant (19, 26).

Extracellular protein toxins have not been identified in *H. somni*. However, high molecular weight fibrillar proteins that bind IgG2 are present on the cell surface of all *H. somni* strains tested except for some preputial isolates (19, 27, 28). This high molecular size, fribillar immunoglubulin binding protein is now referred to as IbpA, and is encoded by the almost 12.3 kb *ibpA* gene (8). Near the C-terminus of IbpA are two direct base pair repeats containing the motif Fic, which has been shown to be cytotoxic for bovine alveolar epithelial cells and phagocytic cells, and can cause the cells to round up and their actin filaments disrupted, which may also inhibit phagocytosis (9). Therefore, *H. somni* may be able to survive within phagocytic cells as a result of compromised cell functions due to cytotoxic effects. Although the isolates lacking IbpA were highly susceptible to killing by BM cells, a few preputial isolates that did produce IbpA (strains 24P, 124P, and 20P) were also killed by BM cells, suggesting there may be factors other than IbpA that contribute to the resistance of *H. somni* to killing by phagocytic cells. To determine if IbpA contributed to bacterial survival through host cell toxicity by the Fic domain (9, 19, 20), semi-purified IbpA was added to the BM cells prior to addition of strain 129Pt, which lacks IbpA and is highly susceptible to intracellular killing. The addition of IbpA to BM cells did enhance the intracellular survival of strain 129Pt, and to a lesser extent other *H. somni* isolates lacking IbpA from the healthy prepuce, and did so in a dose-dependent manner. However, the intracellular killing of 129Pt and other strains was not completely abrogated, and significantly more cells of strain 129Pt were killed by BM cells supplemented with IbpA than strain 2336. Therefore, IbpA may interrupt some essential functions of the phagocytic cells, such as the rearrangement of actin through cytotoxicity (9), but other factors appear to be required to enable *H. somni* to replicate intracellularly.

To more comprehensively examine the role of IbpA in resistance to killing by serum and phagocytic cells, Tn and allelic exchange mutants were tested for intracellular survival. All *ibpA* and other Tn mutants tested were capable of replicating in BM cells. However, all the Tn insertions in the *ibpA* mutants were located near the 5’ end of the *ibpA* gene, which is responsible for immunoglubulin binding through the Fc region. The *N*-terminus of IbpA has homology to the *Bordetella pertussis* filamentous hemagglutinin (Fha), which contributes to adherence (10, 13), and this region is also proposed to be responsible for epithelial cell adherence by *H. somni* (7). These Tn mutants are also deficient in biofilm formation (29), for which the first stage is adherence, and is therefore consisten with a role of the *N*-terminus in bacterial adherence. In contrast, the Fic motifs are located within the DR1/DR2 repeats located near the C-terminus of the protein and were still transcribed in the Tn mutants (data not shown). The Tn insertion may also have caused a frame shift that created a new start codon in the middle of *ibpA*, which is over 12 kbp in size, or the gene remained in frame enabling Fic to be transcribed.

The Fic motifs within DR1/DR2 of the IbpA protein are associated with loss of actin filament function, reduced phagocytosis, and cytotoxicity (7). Therefore, we sought to determine if mutations in specific regions of *ibpA* would negate intracellular survival. Strains that normally lack IbpA or a mutant lacking all of the *ibpA* gene are not cytotoxic (9), as they also lack the cytotoxic Fic motifs within DR1/DR2 (10). However, whether cell cytotoxicity is associated with intracellular survival of *H. somni* has not been examined. Therefore, mutants with in-frame deletions in specific regions of *ibpA* were examined for intracellular survival as well as serum resistance, which is also associated with IbpA (7). All mutants tested with deletions in specific sites throughout *ibpA*, including DR1 and/or DR2 that include the Fic motifs, or a mutant lacking the entire *ibpA* gene replicated in BM cells as effectively as parent strain 2336. Therefore, cytotoxicity of phagocytic cells due to the Fic motifs do not explain the capability of *H. somni* to survive within BM cells.

Bacterial pathogens that can survive within professional phagocytes use one or more mechanisms to avoid intracellular killing, such as: 1) inhibition of phagosome-lysosome fusion and acidification; 2) survival within the phago-lysosome; 3) escape from the phagosome prior to lysosome fusion; 4) killing or lysing of the phagocytic cell (or phagosome or lysosome) before or after phagocytosis (30, 31). We examined co-localization of *H. somni* with intracellular markers to asses traficking of *H. somni* in the phagosome. All *H. somni* strains and mutants tested that could survive or were killed within BM cells co-localized with EEA-1, which is an early endosomal marker, and an early component in phagosome maturation and lysosome fusion (32). However, strain 2336 and other strains that survived within BM cells, whether expressing IbpA or not, failed to acidify phagosomes and did not co-localize with LAMP-2, which is a lysosome-associated membrane protein (32), and an indicator of phagosome-lysosome fusion. The addition of IbpA to BM cells prior to addition of strain 129Pt did not alter acidification of the phagosome or co-localization with LAMP-2, further indicating that Fic toxicity was not responsible for the differences in intracellular traficking noted between strains susceptible or resistant to intracellular killing. The presence of IbpA and DR1/DR2 regions has also been associated with inhibition of uptake by phagocytic cells (7) most likely through disrupting the function of actin filaments (9). However, in these cases either *H. somni* cells expressing IbpA or semi-purified IbpA were added to the phagocytes, incubated for a period of time, and then bacteria (e.g. *S. aureus*) or microparticles added. In these cases, cytotoxicity was likely responsible for reducing subsequent uptake of bacteria or particles. However, there did not appear to be any significant or consistent difference in the uptake of *H. somni* by healthy BM cells, whether the bacteria expressed IbpA or not. Therefore, in addition to the effects of cytotoxicity by Fic on phagocytic cells, most *H. somni* strains also appear to be capable of intracellular survival through inhibition of phagosome-lysosome fusion.

*H. somni* is not unique among mucosal pathogens in being capable of surviving within phagocytic cells. Nontypable *Haemophislus influenzae* is capable of surviving within human THP1 monocytic cells (33), and some strains of *Neisseria gonorrhoeae* and *N. meningitidis* are capable of surviving and thriving within polymorphonuclear leukocytes (34). These bacteria have not been classified as facultative intracellular pathogens because their preferred niiche within the host is not normally within phagocytic cells. Therefore, the term permissive intracellular pathogen may be a more appropriate term for *H. somni* and other typically extracellular mucosal pathogens capable of surviving within professional phagocytes. Whether inhibition of the oxidative burst by *H. somni* (18) is related to their capability to avoid maturation of the phago-lysosome or is an additional mechanism of intracellular survival has yet to be determined, as does the role of intracellular survival in the pathogenesis of *H. somni* diseases.

The expression of IbpA on *H. somni* isolates is also associated with serum resistance (26, 35), as is the structure of the LOS oligosaccharide, and its modification with factors such as sialic acid and phosphorylcholine (36, 37). The association of the entire IbpA protein with serum resistance was confirmed in this study. The isogenic mutant lacking the entire *ibpA* gene was significnatly more serum-sensitive than parent strain 2336 at some dilutions of antiserum. However, the lack of only DR1/DR2 did not increase serum sensivitiy,but in fact made the bacteria more serum-resistant. IbpA was originally identified as an immunoglubulin binding protein, and it is likely that binding of host immunoglobulin through the Fc region may inhibit complement binding and activation (35). Why the lack of DR1/DR2 may enhance serum reistance is unknown, but could be the result of a conformational change in the protein on the cell surface that further blocks complement binding and activation.

## MATERIALS AND METHODS

### Bacterial strains

The *H. somni* strains and mutants used in this study are listed in Table 1. Mutant 2336ΔIbpA1 has almost all of *ibpA* replaced with a kanamycin resistance gene (9), mutant 2336ΔIbpA5 has an in-frame deletion of the 3’-terminal sequence of *ibpA* (AC region; nucleotides 11725-12417 of *ibpA*), mutant 2336ΔIbpA7 has an in-frame deletion of the *ibpA* DR2 sequence (nucleotides 10258-11439 of *ibpA*), mutant 2336ΔIbpA8 has an in-frame deletion of the *ibpA* DR1 sequence (nucleotides 8980-10185 of *ibpA*), mutant 2336ΔIbpA9 has an in-frame deletion of both the DR1 and DR2 sequences (nucleotides 8980-11439 of *ibpA*), and mutant 2336ΔIbpA11 has an in-frame deletion of the *ibpA* R1R2 sequence (nucleotides 6748-8187 of *ibpA*) (8, 38, 39). Preputial isolate 129Pt does not produce IbpA and is not cytotoxic for epithelial cells (19). However, no other differences have been identified in outer membrane proteins examined (40). All strains were grown on Brain Heart Infusion (BHI) agar with 5% sheep blood in 5% CO_2_ overnight from frozen stocks. The colonies were transferred to BHI broth supplemented with 1% yeast extract, 0.1% Trizma base, and 0.01% thiamine monophosphate (TMP) (BHIY-TT) (41), and shaken rapidly (~200 rpm) at 37 °C to mid-log phase.

### Isolation of BPBM and cell lines

Peripheral blood was collected from the jugular vein of Holstein cows into an EDTA-coated tube. Control experiments demonstrated that incubation of *H. somni* strain 129Pt with each preparation of BPBM resulted in very similar killing of strain 129Pt (±<5% difference). The buffy coat layer was separated from the red blood cells and plasma by centrifugation at 1000 × *g* for 30 min at 15 °C, diluted with Hank’s Balanced Salt Solution (HBSS; Life Technologies, Carlsbad, CA) and laid over Ficoll-Paque (Pharmacia, Piscataway, NJ). The BPBM were recovered following centrifugation, as per the manufacturer’s instructions. The viability of the isolated BPBM was greater than 95%, as determined by trypan blue staining, and were grown as confluent monolayers in RPMI-1640 medium supplemented with 10% fetal bovine serum. A bovine monocytic cell line (BM) derived from blood monocytes of a 6-year-old Guernsey cow was originally obtained from Dr. John Dame, University of Florida, and provided by Dr. David Lindsay, Virginia Tech. The cells were grown in RPMI-1640 medium supplemented with 10% heat-inactivated fetal bovine serum and 2 mM L-glutamine in 5% CO_2_ at 37 °C (42). The cells were cultured in wells of 6-well plates at 3×10^5^/well (in 5 ml), allowed to adhere for 24 h, and the monolayers were washed 3 times with phosphate buffered saline, pH 7.2 (PBS) before adding IbpA or bacteria. Other cell lines tested in the same manner were mouse J774A.1, human THP, and bovine FBM-117 (43).

### Construction of *ibpA* allelic exchange and transposon (Tn) mutants

Seven mutants (#9, #3, #13, #91, #27, #23 and #137) with a Tn insertion within *ibpA* were selected from a bank of mutants made with EZ::Tn5™<KAN-2>Tnp Transposome™ (Epicentre, Chicago, IL) as previously described (29). The Tn insertion site was confirmed by sequencing the ends of the Tn and the flanking chromosomal region. Deletion mutant 2336ΔIbpA1 was made by replacement of essentially all of the *ibpA* with a kanamycin resistance gene (Kn^R^) (9). Mutants 2336ΔIbpA5, 2336ΔIbpA7, 2336ΔIbpA8, 2336ΔIbpA9, and 2336ΔIbpA11 were made by in-frame deletions using a temperature-sensitivite plasmid (38, 39).

### Preparation of IbpA

The IbpA protein secreted into the culture medium was concentrated as previously described with minor modifications (13). Briefly, *H. somni* strain 2336 was grown in 300 ml of BHIY-TT to mid-log phase. The bacteria were removed by centrifugation at 8000 × *g* for 10 min. The supernatant was filtered through a 0.2-μM filter to remove any residual bacteria and then concentrated to 15 ml (20-fold) through a 10,000 MW centriprep centrifugation filter unit (Millipore, Billerica, MA) by centrifugation at 4,000 × *g* for 30 min at 4°C. The retentate was used as a source of concentrated IbpA in the phagocytosis assay. The purity of IbpA could not be determined because the protein is very large and consists of subunits or aggregates of 76, 120, 270, and 370 kDa (7). In Western blots more than 20 proteins from 76 kDa to 370 kDa are present in IbpA-positive strains, but none are present in IbpA-negative strains, indicating this procedure results in a sample consisting of predominately IbpA (19).

### Phagocytosis assay

Bacteria in mid-log phase were co-incubated with BPBM or BM monocytic cells for 1 h at a 100:1 multiplicity of infection (bacteria:monocytes). The monocytes were then incubated with 50 μg/ml of gentamicin for 30 min to kill extracellular bacteria, washed three times with PBS, and incubated at 37 °C. After incubation for 0 h (1 h after addition of bacteria and 30 min after addition of gentamicin), 24 h, 48 h, or 72 h the monocytes were lysed with distilled water, neutralized with 2X PBS, and the lysate cultured onto BHI blood agar to determine the number of viable intracellular bacteria. The uptake of *H. somni* by the monocytes was determined at time 0, following lysis of the monocytes and inoculation to BHI blood agar. The capability of IbpA to protect bacteria from intracellular killing by the monocytes was determined by co-incubation of the cells with 20-fold concentrated IbpA (3.6 mg/ml), or a sample diluted to represent a 4-fold concentration, from culture supernatant for 2 h before co-incubation with the bacteria.

### Confocal Microscopy

To determine the intracellular trafficking of the bacteria in BM cells, two groups of BM cells were incubated with bacteria for different time periods. In group 1 BM cells were incubated with the bacteria for 30 min at 37°C, the culture medium with nonadherent bacteria was removed by washing the monolayer gently three times with PBS, and the medium was replaced with 3% paraformaldehyde for 10 min to fix the cells. Another group of BM cells was also incubated with the bacteria as above, but after removal of the culture medium, fresh medium was added and incubation was continued for 3 hr at 37°C. The BM cells were then washed and fixed as above for 10 min. The cells were then permeabilized with 0.1% saponin solution containing 1% bovine serum albumin (BSA) and 5% goat serum for 30 min. The intracellular *H. somni* and phgosomal/lysosomal markers were labeled with rabbit antibodies to *H. somni*, or EEA-1 or LAMP-2 and visualized with goat anti-rabbit antibodies conjugated with Alexa Fluor^®^ 488 (*H. somni*) or Alexa Fluor^®^ 546 (markers) (Life Technologies, Carlsbad, CA) by confocal microscopy (Zeiss LSM 510 META; Carl Zeiss, Thornwood N.Y.). The nucleus of the monocytes was visualized with DAPI (Life Technologies, Carlsbad, CA). To determine phagosome acidification, the monocytes infected with *H. somni* were incubated with Lysotracker (Molecular Probes, Eugene, OR) for 1h, following the manufacturer’s instructions. The cells were then fixed as described above. To determine the percentage of *H. somni* co-localized with the markers, the percent of *H. somni* co-localized with each marker was determined from the number of *H. somni* co-localized with either EEA-1 or LAMP-2 divided by the total number of *H. somni* cells. The markers and intracellular *H. somni* were identified by SpotFinder Z and counted manually in 5 fields, each containing at least 4-5 monocytes (44, 45). Quantification of each marker, including LysoTracker, was determined using the software imageJ (https://imagej.nih.gov/ij/download.html) and MicrobeTracker (http://www.microbetracker.org/), which analyze the percentage of fluorescent dots and fluorescence signals, respectively.

### Statistical analyses

Two-tailed P values were calculated using the unpaired *t* test. A P value <0.05 was considered significant. Statistical analyses were determined using InStat 3 software (GraphPad Software, Inc., La Jolla, CA).

## ACKNOWLEDGEMENTS

We would like to tank Dr. David Lindsay for providing the BM macrophage cell line, Dr. Lynette Corbeil and Vivian Fussing for providing bacterial strains, and Poorna Goswami, Angelea Sadaat, and Gillian Rodgers for excellent technical assistance. This work was supported by USDA-NIFA grant 2013-67015-21314 to TJI and by HATCH funds from the Virginia-Maryland College of Veterinary Medicine.

